# Configuring a federated network of real-world patient health data for multimodal deep learning prediction of health outcomes

**DOI:** 10.1101/2021.10.30.466612

**Authors:** Christian Haudenschild, Louis Vaickus, Joshua Levy

## Abstract

Vast quantities of electronic patient medical data are currently being collated and processed in large federated data repositories. For instance, TriNetX, Inc., a global health research network, has access to more than 300 million patients, sourced from healthcare organizations, biopharmaceutical companies, and contract research organizations. As such, pipelines that are able to algorithmically extract huge quantities of patient data from multiple modalities present opportunities to leverage machine learning and deep learning approaches with the possibility of generating actionable insight. In this work, we present a modular, semi-automated end-to-end machine and deep learning pipeline designed to interface with a federated network of structured patient data. This proof-of-concept pipeline is disease-agnostic, scalable, and requires little domain expertise and manual feature engineering in order to quickly produce results for the case of a user-defined binary outcome event. We demonstrate the pipeline’s efficacy with three different disease workflows, with high discriminatory power achieved in all cases.

## 1 INTRODUCTION

In recent years, there has been an explosion in the amount of collected structured and unstructured clinical data in the form of Electronic Health Records (EHR) and Electronic Medical Records (EMR) [22]. Concurrent advancements in machine learning (a set of computational heuristics that seek to identify nonlinear interactions and transformations) have simultaneously made big data analysis more tractable. Deep learning methods operate on data using artificial neural networks. These complex models are capable of recognizing patterns in large-scale, multidimensional data [8], and have seen successful applications across multiple health-care domains, such as clinical imaging (particularly in pathology and radiology image recognition and segmentation), genomics, and more recently, wearable health monitoring devices [18].

In this study, we consider EHR data which is inherently multimodal and is comprised of structured components (e.g. diagnoses, procedures, laboratory values, vital signs, and demographic information), and unstructured components (e.g. medical images and clinical free text). The structured data are encoded using a variety of medical classification lists. Diagnoses are most commonly encoded with International Classification of Disease (ICD) codes, procedures are represented by Current Procedural Terminology (CPT)/ Healthcare Common Procedure Coding System (HCPCS) or ICD Procedure Coding System (ICD-PCS), and laboratory and vital values with Logical Observation Identifiers Names and Codes (LOINC). Models integrating this information across large federated datasets that are aggregated across many institutions around the United States may produce better discriminatory results versus training within a single institution but are difficult to train because they require standardization and completeness across these training datasets as well as privileged access to highly sensitive data. In addition, mapping these data manually to a consistent ontology is labor-intensive, and periodic revisions necessitate refactoring of existing pipelines and retraining personnel (e.g., ICD-9 to ICD-10 transition in late 2015) [25].

Another barrier to entry for these emerging technologies is the lack of interpretability of deep learning approaches versus standard algorithms deployed in the healthcare fields, such as logistic regression. The latter involves sensible curation of clinically relevant covariates, while the former involves an approach that freely uses all available covariates while largely obfuscating which factors the model found to be clinically-relevant. This so called “Black Box” interpretability problem makes clinician acceptance and use of any targetable insights derived therein more difficult to implement. However, deep learning approaches have significant advantages over simpler models trained on human curated data in that they incorporate feature extraction into the learning process of the network instead of the manual specification of covariates and thus consider associations a human observer might dismiss or fail to recognize. Examples specific to the EMR/EHR deep learning field include the stacked autoencoder-based DeepPatient, proposed by Miotto et. al. [19], the concept embedding-based Med2Vec, proposed by Choi et. al. [2, 3, 5], and Rajkomar et. al.’s method based on the Fast Healthcare Interoperability Resources (FHIR) format [23].

Another significant issue in training multi-institutional deep learning models on patient data is the Protected Health Information (PHI) contained in EHR data. Hospitals are understandably hesitant to disseminate HIPAA-protected data outside of their own firewalls due to the risk and penalties of data leakage. Indeed, most hospitals are so cognizant of this risk as to limit the access of practitioners within their own institutions to only the patient information that is necessary for a clinician to perform their duties. A data protection scheme, such as federated learning, where patient data is encrypted and processed behind the firewall of the originating institution before being used in a federated learning model is necessary for any model requiring broad access to protected patient data [24, 29].

In this work, we use a data repository of EHR data (TriNetX’s Federated Network) that has been aggregated and homogenized from multiple member institutions, which include healthcare organizations, biopharmaceutical companies, and contract research organizations [28]. These institutions pool and share data to recruit patients for clinical trials and conduct research using the TriNetX platform. In contrast to the aforementioned federated learning platforms, from which noisy/heterogenous data can potentially corrupt models trained across institutions, curation of pooled data resources in a secure centralized, collaborative platform allows for greater standardization of resources, improving research quality. We provide further support for the utilization of computational pipelines that are built on EMR data that generalize to a range of binary outcomes. Our framework tackles complications associated with dataset completeness, while demonstrating the capabilities of data extraction from large federated data networks and integration into deep learning workflows.

## 2 MATERIALS AND METHODS

### 2.1 Data Query Mechanism

We developed an auto-querying mechanism to extract patient data from TriNetX’s Dataworks network, a subset of their full network containing approximately 48 million patients from 39 healthcare organizations, that has been deidentified and approved for research use [7]. The auto-querying mechanism constructs a training cohort through user-specification of a binary-outcome defined by one or more ICD-9/10 codes or, in the case of a histological outcome, ICD-O codes (presence of the code indicates a “positive” patient). The pipeline automatically generates a matched negative population through automatic, greedy and exact identification of age- and sex-matched controls in a general patient pool who lack the specified ICD codes. Additional temporal constraints may be placed to ensure matched patients had unrelated diagnoses recorded on the same day as the positive patient’s index date. Using the auto-querying mechanism, the user can then specify one or more health metrics that they wish to obtain from those populations for processing and analysis, including: diagnoses (including those from the tumor registry), laboratory values, vitals, procedures, and demographic data. The software automatically generates statistical reports to ensure that the age and sex distributions are matched between cohorts.

### 2.2 Data Preprocessing

Once the data has been acquired (**Figure 1A**), the data is split into two separate preprocessing pipelines devoted to code-based categorical data (e.g., diagnoses and procedures; ICD-9, ICD-10, ICD-9-CM, ICD-10-PCS and CPT codes) and value-based data (e.g., laboratory values, vital values, and demographic data).

**Figure 1.**
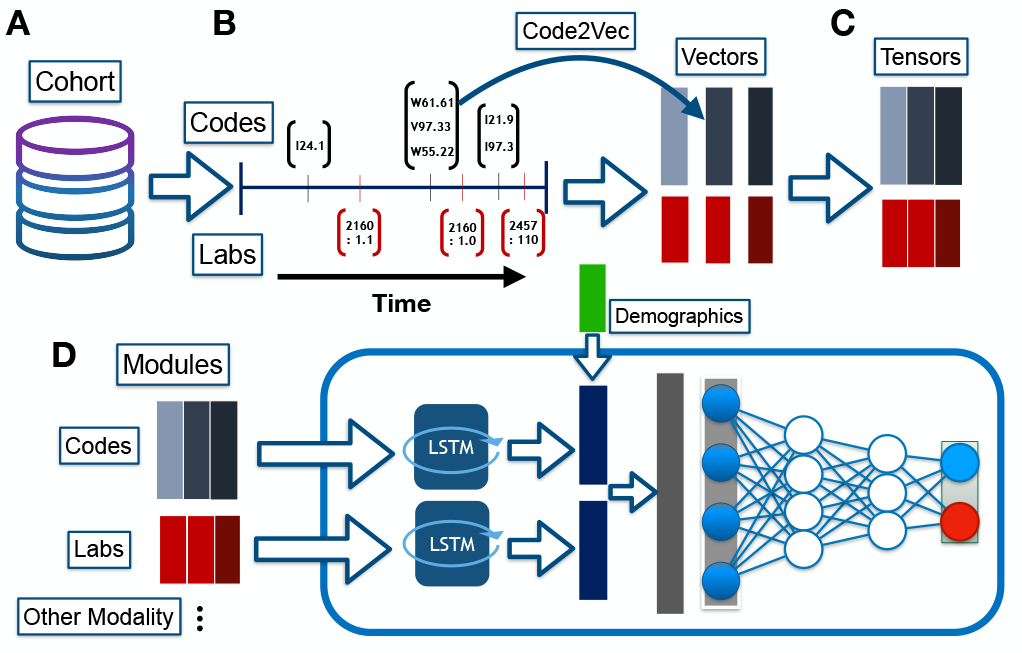
Graphical overview of modeling approach. A) Cohort is generated from federated data network; B) Codes and laboratory measurements (LOINC) extracted over time (referenced by index date) and are converted to their vectorized representations for the timesteps; C) Vectors across time are concatenated to form sequences of information (tensors), which serve as input for D) the Modular LSTM, which processes the temporal sequences across multiple modalities, combines this information with demographics and further operates on the data using multi-layer perceptrons to output a positive/negative diagnosis

#### 2.2.1. Code2Vec to encode diagnostic and procedural information based on neighboring codes

Prior to encoding diagnostic and procedural codes Contextual vectorized representations of diagnosis and procedure codes (i.e., codes are assigned to vectors which are compared to one another using vector arithmetic), enumerated above, are created using the Word2Vec algorithm [1, 15] (**Figure 1B**; **Appendix Table 1**). The dimensionality of the representation is user-definable, but a 100-dimensional embedding is recommended for an optimal balance between computational cost and a capture of the entirety of the feature space. Prior to encoding these codes, ICD-9 codes are mapped to ICD-10 using a standardized mapping dictionary [21]. The Word2Vec dictionary is built using codes from both the negative and positive population cohorts to ensure unknown codes are recognized.

#### 2.2.2. Value-based data preprocessing

Primary value-based data include laboratory (clinical chemistry, hematology, microbiology, etc.) and vitals data, encoded with LOINC codes [14] (**Figure 1B**). Uncommon laboratory measurements are excluded via frequency thresholding, where codes must be present across a user-specified fraction of the population. Sex and year of birth are imputed with the most common sex and combined mean of year of birth (year of birth as proxy for age).

#### 2.2.3. Removal of endogenous information via temporal filtering

A temporal filtration system prunes the code and value-based record information from a user-defined number of days before the event (patients in the negative population also have an index event date; selection of an index date is common practice for selection of negative controls, where index dates are matched between the positive/negative cohorts) [12]. The selection of the temporal filter window is largely dependent on the disease being studied, patient numbers, types of data included, and other clinical considerations particular to the condition to ensure meaningful prospective prediction and avoid endogeneity (e.g., analysis of a seasonal influenza).

#### 2.2.3. Final data representations

The final dataset is represented using datatypes that reflect either (**Figure 1C**):

1. Temporal sequences (sequence of codes or lab measurements)– Diagnostic and process/procedure codes are converted into their word2vec vectors, which are tagged and then ordered by date of assignment. If multiple codes occur on the same day, the vectorized representations are averaged to summarize the day. The resulting vectors are ordered temporally, with each diagnosis/procedure vector representing a discrete patient encounter. Each patient is assigned a sequence of Code2Vec vectors (*n* Code2Vec dimensions by *t* timepoints per patient). Value-based records are represented by a multivariate time-series. For instance, each type of clinical lab measurement (e.g., levels of hemoglobin, sodium, etc.) serves as an individual time series. Measurements are averaged and mean-imputed for each day (sequence of multiple lab measurements, *m* lab measurements by *t* timepoints per patient).
2. Static representation– instead of representing codes and lab measurements as sequences, the code and value-based measures can be averaged across the time period. Fixed demographic clinical covariates serve as additional predictors.

The user has the option to remove patients that do not utilize all of the available datatypes (absence of codes, lab measurements and demographic information).

### 2.3 Modeling Approaches

Our framework operates on static data representations with the following machine learning models: Logistic Regression (L2-regularized), Multi-Layer Perceptrons (MLP; trained with the Adam optimizer with a learning rate of 1e-3, trained until the plateau of the validation loss or for 200 epochs), Random Forest Classifier (ensemble method; decision splits decided based on Gini impurity). Finally, a 2-layer bi-directional Long Short-Term Memory (bi-LSTM) network (which retains information from intermediate hidden states of the model) may be employed to learn temporal dependency between the code and value-based measurements. The final hidden state is concatenated with representations of the other data modalities (e.g., combining codes with lab measurements and demographics) to yield an ultimate data representation. The final data representation is passed through a final MLP, where a sigmoidal activation function is applied to obtain class probabilities. We denote this approach as the modular LSTM (mLSTM) approach, as other EHR modalities represent interchangeable “modules” (**Figure 1D**). The default training scheme utilizes the Adam optimizer with minibatches and a cross-entropy loss function. Dataset bias from imbalanced class distributions were ameliorated using reweighting of the model objective function, oversampling of the low-frequency class, or a sensitivity analysis of the receiver operating curve (ROC).

### 2.4 Model Interpretation

#### 2.4.1. Interrogation of Semantic Similarity between Code2Vec Vectors with T-SNE

Given that the Code2Vec vectors are being modeled sequentially using a Bi-LSTM to make disease predictions, it is expected codes can be contextualized by the type of malignancy and potential co-morbidities with other codes since co-morbidities co-occur (are both present despite being placed in different parts of the ICD code hierarchy) within similar time windows. We verified that the Code2Vec model had registered the correct semantic information through inspection of T-Stochastic Neighborhood Embedding (T-SNE) plots, which generate 2-D visualizations of the 100-dimensional Code2Vec vectors by projecting high dimensional data onto a manifold which preserves semantic distances between codes [11]. Codes were colored by type of malignancy and in the results section, we comment on the potential similarity between co-morbidities.

#### 2.4.2. Integrated Gradients for Codes and Lab Measurements

Integrated Gradients is a post-hoc model explanation technique which can assign an importance score to any predictor of the model for their relevance for the prediction. We used the Integrated Gradients technique to establish which codes were important for prediction of the disease at each time step, per patient by backpropagating information of prediction back to the original input (one score per Code2Vec dimension, per timepoint). The overall importance for a given timestep was given by averaging the score across all Code2Vec dimensions across the timestep (alternatively, the absolute value of the integrated gradient score may be taken to remove relationship directionality). The importances of the multivariate clinical lab measurement time series were calculated similarly, where instead the output remains unmanipulated, allowing for the interrogation of the importance of a specific lab measurement across time.

### 2.5 Experimental Design

We tested our pipeline on three separate patient cohorts corresponding to the following conditions: pancreatic cancer (PC), colorectal cancer (CRC), and non-alcoholic steatohepatitis (NASH). These patient cohorts were defined on the basis of a recorded ICD-9/ICD-10 diagnosis code corresponding to each of the three conditions on their medical record. Our framework split the datasets into training, validation, and testing sets, with the default splits being 80%, 10%, and 10% respectively before testing on each type of classifier. Predictive performance was measured using the area under the receiver operating curve (AUROC; estimate of the probability that a randomly selected case has a greater predictive probability than a randomly selected control), balanced accuracy and F1-Score (geometric mean of model precision and recall) measures.

## 3 RESULTS AND DISCUSSION

### 3.1 Results

#### 3.1.1. Age, sex matching for harmonized data pull

Population characteristics such as age and sex exhibited similar distributions in positive and negative testing populations (**Table 1**).

**Table 1:** Demographic characteristics of the training populations. Negative populations were age-, sex-, and temporally matched to the positive population in a greedy fashion, and a ten-day filtration was used for all data types. The cohort selection process is described in detail in the pipeline description section. For pancreatic cancer, a single negative match per positive patient was specified. For colorectal cancer and non-alcoholic steatohepatitis (NASH), a maximum of two negative patients were matched to a positive patient, in order to generate approximately balanced population numbers.

|  | Pancreatic cancer |  | Colorectal cancer |  | NASH |  |
| --- | --- | --- | --- | --- | --- | --- |
|  | + | - | + | - | + | - |
|  | n = 10150 (53.7%) | n = 8746 (46.3 %) | n = 21485 (48.1%) | n = 23223 (51.94%) | n = 11178 (51.5%) | n = 10508 (48.5%) |
| **Year of birth** |  |  |  |  |  |  |
| Mean (SD) | 1948.5 (13.0) | 1948.5 (14.0) | 1948.6 (13.3) | 1950.6 (15.7) | 1958.4 (13.5) | 1957.9 (16.0) |
| Median | 1947 | 1947 | 1947 | 1948 | 1956 | 1956 |
| **Sex (N, %)** |  |  |  |  |  |  |
| Male | 5183 (51.1) | 4531 (51.8) | 10415 (48.5) | 11810 (50.9) | 4795 (42.9) | 4803 (45.7) |
| Female | 4967 (48.9) | 4215 (48.2) | 11070 (51.5) | 11413 (49.2) | 6383 (57.1) | 5705 (54.3) |

#### 3.1.2. Predictions Across Disease Subtypes

We fit four modeling approaches to the federated dataset for the tasks of predicting the likelihood of pancreatic cancer, colorectal cancer and NASH on a multimodal dataset (in this instance, diagnostic codes, laboratory data and demographic characteristics). In **Table 2**, we present fit statistics on the held-out test sets. Prediction of pancreatic cancer across all classifiers yielded AUROC scores of 0.78 – 0.87, while colorectal cancer was the least characterizable, with models achieving 0.71 – 0.82 AUROC. Finally, presence or absence of NASH was the most characterizable, with the various model types achieving AUROC scores of 0.84 – 0.91 (**Table 2**).

#### 3.1.3. Performance Comparison Across Models

Overall, the mLSTM model outperformed all other modeling approaches for all three datasets (**Figure 2**) across all metrics, with AUROC of 0.871 (PC), 0.724 (CRC), and 0.841 (NASH). The difference is most notable in the case of pancreatic cancer prediction but is still present in the colorectal cancer and NASH workflows. Generally, the MLP and random forest classifier models underperformed compared to the mLSTM and logistic regression. Deep learning models exhibited higher recall/sensitivity at a standard threshold than the logistic and random forest models, though in the case of the MLP, this sensitivity comes with an increase in the false positive rate (Table 2). Addition of data types, including laboratory values and demographic data, to diagnostic information results in an increased AUROC over diagnosis information alone (**Table 3**).

#### 3.1.4. Code2Vec embeddings capture disease comorbidities

Inspection of the latent Word2Vec embeddings of diagnostic codes demonstrated high clustering of diagnostic and procedural codes by malignancy types and common comorbidities (**Figure 3**).

**Table 2:** Test set performance on the three disease workflows. Processed patient data was split into a training, validation, and testing set with sizes 80%, 10%, and 10% (random seed = 10). All long short-term memory (LSTM) models took in temporally-processed versions of diagnosis and laboratory data, with age/sex data represented statistically. Models were bilayer and trained over 30 epochs. No dropout regularization was used. Logistic regression used L2 regularization (C=1), and was fitted to the static representations of the diagnostic/lab/demographic data (detailed in the preprocessing section). The multi-layer perceptron (MLP) models were 3 layers deep, with 100 neurons per layer. The Random Forest Classifiers (RFC) were comprised of 10 estimators each, with no constraints on individual tree depth.

|  | Pancreatic cancer<br>(Test set n = 1890) |  |  |  | Colorectal cancer<br>(Test set n = 4471) |  |  |  | NASH<br>(Test set n = 2169) |  |  |  |
| --- | --- | --- | --- | --- | --- | --- | --- | --- | --- | --- | --- | --- |
|  | mLSTM | Logistic | MLP | RFC | mLSTM | Logistic | MLP | RFC | mLSTM | Logistic | MLP | RFC |
| **Accuracy** | 79.58% | 73.39% | 72.70% | 69.63% | 74.52% | 71.97% | 70.32% | 66.23% | 84.32% | 79.85% | 79.30% | 78.56% |
| **Bal. Acc.** | 79.54% | 73.58% | 72.34% | 69.90% | 74.34% | 71.57% | 70.30% | 65.52% | 84.47% | 79.88% | 79.26% | 78.82% |
| **AUROC** | 0.871 | 0.810 | 0.800 | 0.777 | 0.817 | 0.782 | 0.768 | 0.710 | 0.911 | 0.878 | 0.881 | 0.864 |
| **Recall** | 0.801 | 0.704 | 0.783 | 0.654 | 0.713 | 0.651 | 0.700 | 0.540 | 0.803 | 0.790 | 0.804 | 0.711 |
| **Specificity** | 0.789 | 0.768 | 0.663 | 0.744 | 0.773 | 0.781 | 0.706 | 0.770 | 0.886 | 0.807 | 0.781 | 0.865 |
| **Precision** | 0.811 | 0.774 | 0.724 | 0.743 | 0.735 | 0.724 | 0.678 | 0.675 | 0.883 | 0.814 | 0.797 | 0.850 |
| **F1 Score** | 0.806 | 0.737 | 0.753 | 0.695 | 0.724 | 0.685 | 0.689 | 0.600 | 0.841 | 0.802 | 0.801 | 0.774 |

**Table 3:**
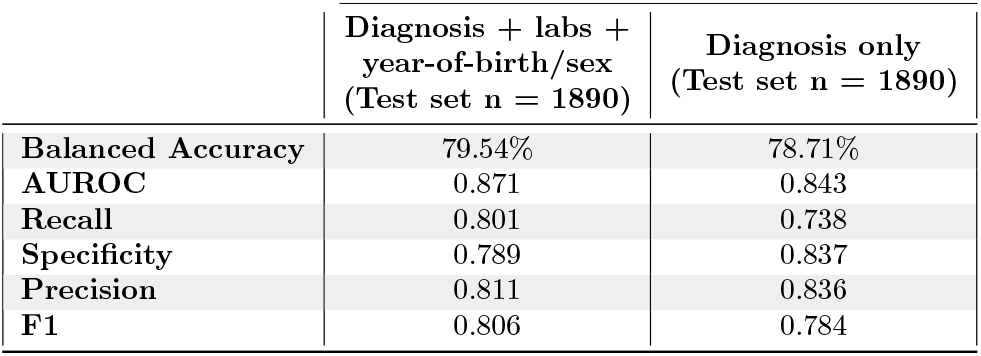
Example of the performance increase afforded by the addition of more data types in the pancreatic cancer workflow. Two temporal model networks were trained under identical conditions, with one model including laboratory values and the other two featuring demographic characteristics, age and sex.

|  | Diagnosis + labs +<br>year-of-birth/sex<br>(Test set n = 1890) | Diagnosis only<br>(Test set n = 1890) |
| --- | --- | --- |
| Balanced Accuracy | 79.54% | 78.71% |
| AUROC | 0.871 | 0.843 |
| Recall | 0.801 | 0.738 |
| Specificity | 0.789 | 0.837 |
| Precision | 0.811 | 0.836 |
| F1 | 0.806 | 0.784 |

**Figure 2.**
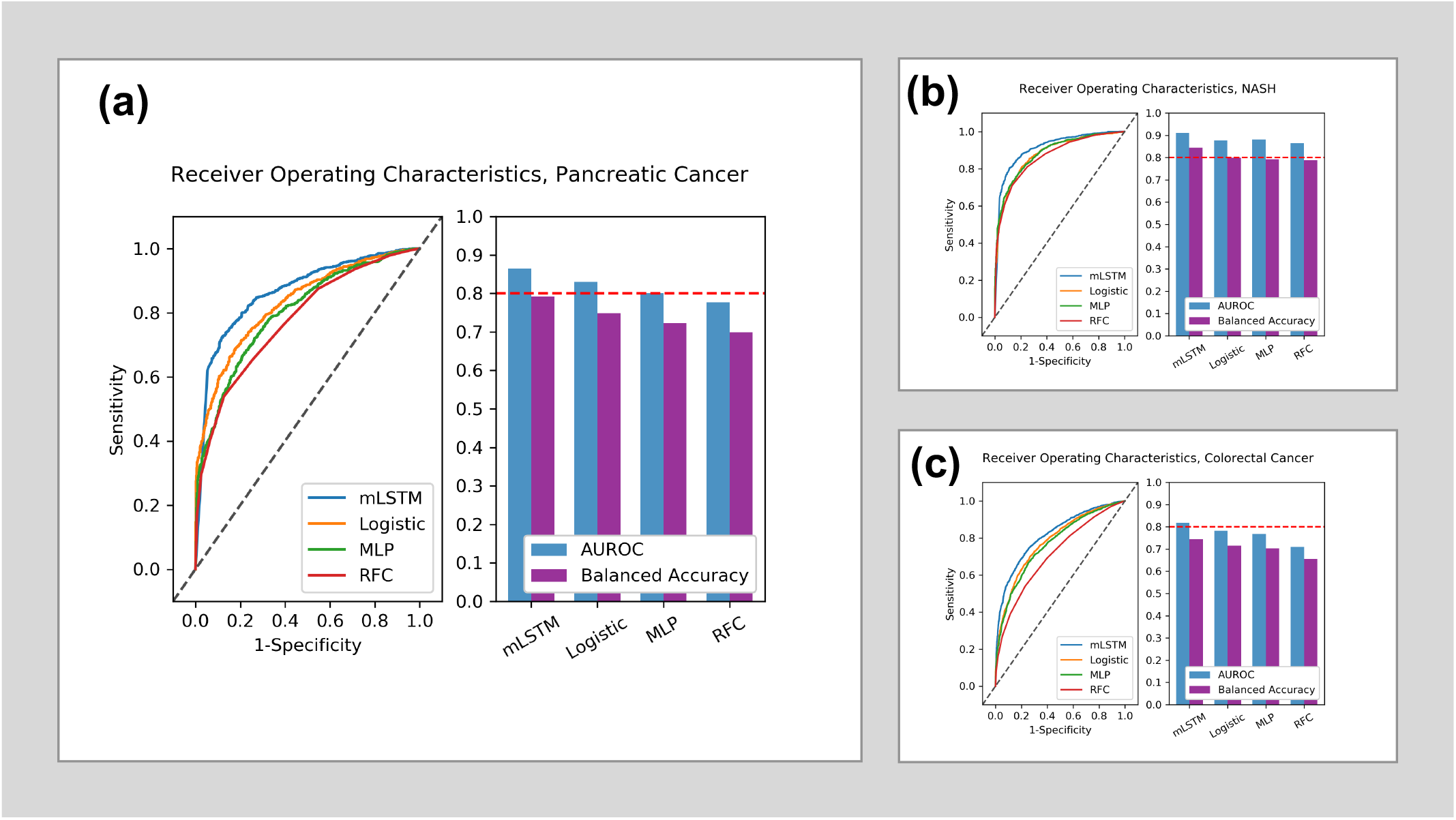
Receiver operating characteristics for (a) pancreatic cancer, (b) Non-alcoholic steatohepatitis (NASH), and (c) colorectal cancer. A reference line at y = 0.8 (dotted red) is added for clarity. All models included diagnosis data, laboratory data, and demographic (age/sex) data. Sample sizes are given in Table 1. A temporal filtration of 10 days from the index event was specified for both diagnostic and laboratory data. Laboratory values were included with default threshold of 0.5, resulting in 27 laboratory values considered for all diseases. Default training settings were used for all models (see pipeline description). Balanced accuracy was calculated by macro-averaging the proportion correct for each class individually. Abbreviations: Area under the Receiver Operating Characteristic Curve (AUROC), modular long short-term memory network (mLSTM), multi-layer perceptron (MLP), random forest classifier (RFC).

**Figure 3:**
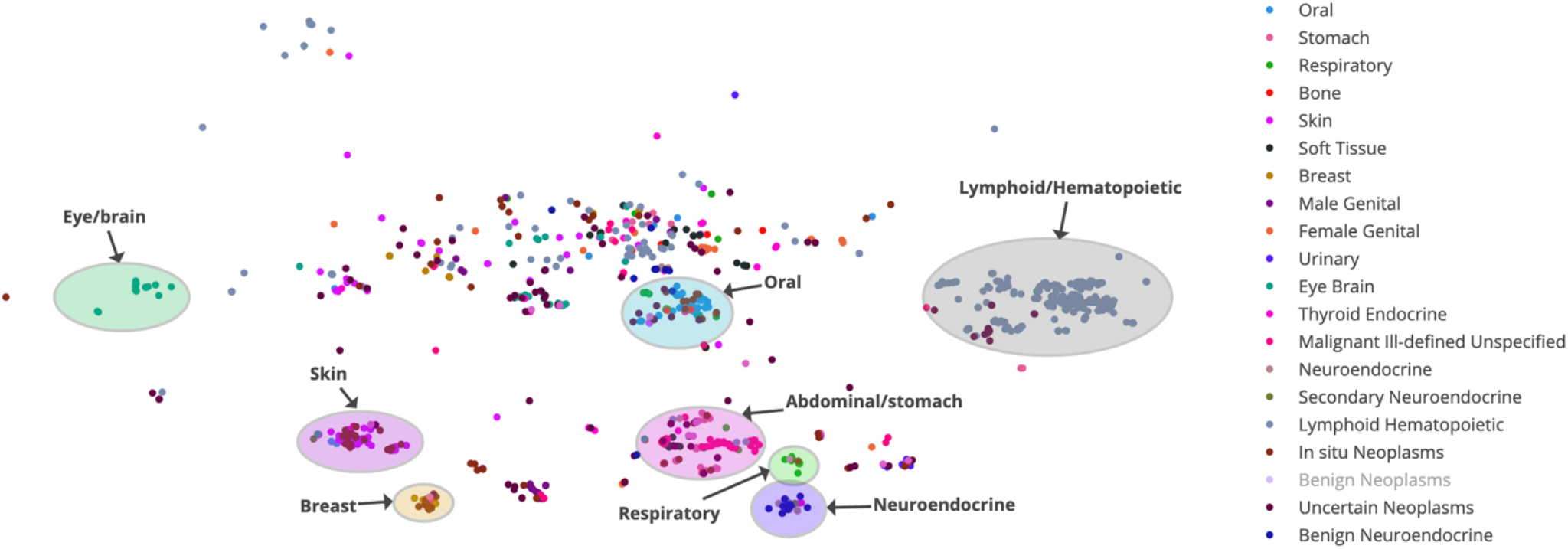
Learned embeddings of malignancy codes with the Word2Vec algorithm on a patient population of 60,000 randomly selected patients from the TriNetX Dataworks network. Each point represents a single ICD-10 code. The original embedding dimensionality was 100, which has been dimensionally reduced through t-distributed stochastic neighbor embedding (t-SNE)^38^. For the purposes of this visualization, only malignancy codes (ICD families beginning with ‘C’ or ‘D’) were included and benign neoplasms were excluded. Codes are categorized and colored by malignancy type, and general groupings of ‘comorbidity spaces’ are labeled.

#### 3.1.5. Temporal importance of code assignments via Integrated Gradients

After applying Integrated Gradients to two randomly selected NASH cases from the test set, we extracted important codes which are likely co-morbid with the underlying condition prior to the final diagnosis. For instance, for the first NASH patient (**Figure 4A**), prior to diagnosis, we noted that abdominal pain (ICD 789) accompanied a diagnosis of other specific liver diseases (ICD K76.89), though NASH was not officially diagnosed until later on. In addition, the patient previously diagnosed with type 2 diabetes (ICD E11.9), a comorbidity that is strongly associated with NASH [6]. The second NASH patient (**Figure 4B**) also had a previous diagnosis of Hepatitis C (ICD 070.54/070.70) and diagnosis of a metabolic syndrome (ICD E88.89), amongst many other co-morbidities, prior to the NASH diagnosis.

**Figure 4:**
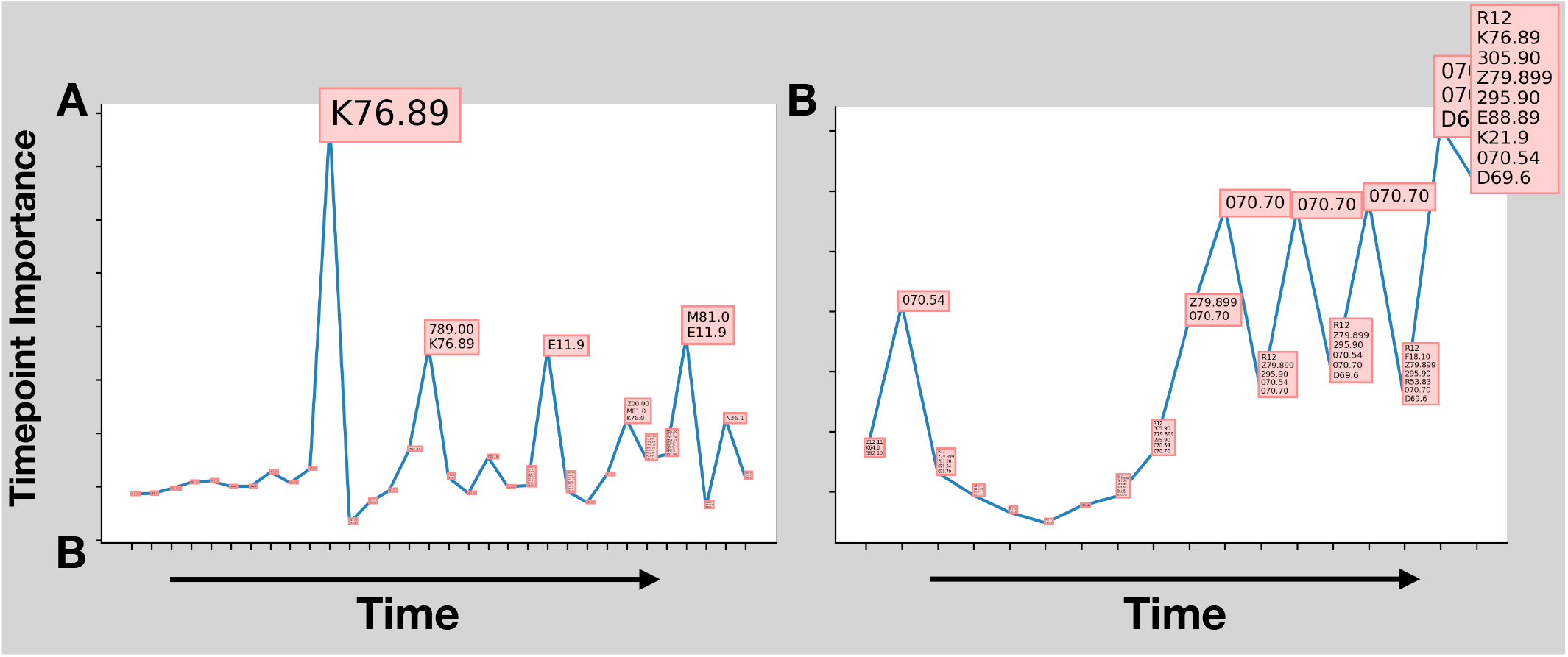
Codes deemed important using the Integrated Gradients algorithm for two select cases (A-B) for Non-alcoholic steatohepatitis. The x-axis indicates the time step since the index event, while the y-axis captures the predictor importance; text containing the ICD codes assigned that day are plotted over their x,y coordinates and sized by their importance

### 3.2 Discussion

Many approaches have been taken with regards to machine/deep learning modeling of electronic patient data [26], for a variety of use cases. Commonly, automated feature representation and outcome prediction are combined into a single workflow, as is the case with our approach. However, our workflow additionally considers processing dilemmas that are characteristic of large healthcare datasets. In this study, we demonstrated successful integration of a harmonized dataset from a large federated network into deep learning modeling workflows which were executed across three separate and meaningful datasets.

TriNetX consists of a large federated network of heterogeneous healthcare entities. The data within TriNetX is ingested from these disparate institutions and companies, quality controlled and standardized, and updated on a regular basis, which provides significant advantages over a static, single institution-sourced datasets. Using a federated, nationally representative dataset allows researchers to train models on diverse patient populations and reduce institution-specific idiosyncrasies and biases. Data volume is a well-established limitation in the deep learning space, with many healthcare datasets being of insufficient size to train the complex, high-parameter models found in deep learning, especially given the judicious amount of preprocessing and cleaning that healthcare data requires. Federated data provides the scope of data necessary to make deep learning a viable modeling method.

Of immense value to this work, we sought to predict whether a patient has a malignancy or other serious disease by integrating temporal information across multiple sources (e.g., codes, labs). While such temporal data is notoriously difficult to work with, given data quality limitations which will be elaborated on, results indicate the potential to deploy these technologies prospectively to shunt a patient into the pipeline earlier than they otherwise would be.

#### 3.2.1. Model Performance

Though direct comparison of models is difficult, for the task of pancreatic cancer prediction, our method performs comparably to the deep-learning based approach proposed by Muhammad et. al. [20] on both EMR and survey data with 18 personal health features. Our pipeline does not utilize survey data and only offers a limited view into behavioral characteristics (i.e., we did not include self-reported measures, which may be unreliable). The high prevalence of pancreatic cancer comorbidities and risk factors and the lack of pathognomonic symptoms makes selection of a patient for further screening challenging. Unsurprisingly, NASH, an inflammatory disease that is the most common cause of end-stage liver disease in the western world, is associated with disease symptomology that is more likely to be captured in clinical records and laboratory tests and as such prediction of NASH outperformed the other two datasets.

NASH is characterized by a disease course that progresses through well-defined stages, beginning with hepatic steatosis and non-alcoholic fatty liver disease before progressing to NASH, with those early diseases frequently presenting with physical exam findings and hepatobiliary lab abnormalities. Additionally, NASH is strongly associated with metabolic disease as well as other comorbidities [13]. The NASH patient case study presented with multiple abdominal- and hepatic-related diagnoses prior to their NASH diagnosis. The course of this patient is consistent with the natural history of NASH, with a patient history of steadily progressive hepatic disease before end-stage liver disease.

#### 3.2.2. Study Limitations

There are several limitations in this study. As our selection of training populations is done in a case-control fashion, our method is subject to all of the biases and limitations that such a study design entails [27]. Matching can ameliorate these biases to some extent. Temporal matching between patients in positive and negative populations by an index event can combat data leakage by avoiding endogeneity and partially account for biases that occur due to changes in diagnostic practice over time (e.g. the shift from ICD-9 to ICD-10 in 2015, which may have altered the presence of certain codes during that time period).

Given the limited transparency in the current version of the pipeline, it is difficult to evaluate whether some unrecognized biases exist between the populations that are artificially inflating the discriminatory capacity of the classification models. Health record data are intended to be a reasonable accounting of the time at which a diagnosis was made. However, diagnosis delays and administrative/insurance issues may prevent a diagnosis or a lab from being recorded on the day it was administered. The temporal filtration of a certain number of days prior to the index date of a patient’s diagnosis of interest was designed to account for data leakage due to recording issues.

Federated networks are subject to data loss during the process of data ingestion and standardization from the various input sources of the network. The internal validation at TriNetX is designed to account for this loss, but some degree of incorrect data collation is unavoidable. However, the large quantity of data in such federated networks serves to reduce the effects of minor errors that arise during the ingestion process, barring a systematic error in data intake.

Addition of model-agnostic post-hoc explainability techniques, such as the Shapley Additive Explanation (SHAP) algorithm [10] and attention, will serve as further validation of the pipeline. Applying self-attention across the modalities may lower the contribution from a “noisy” or “missing” dataset, improving prediction robustness. The Code2Vec results demonstrated promising results for contextual representations of ICD codes and may be further improved by adoption of transformer architectures.

A major strength of our method is the data it is built upon, with the TriNetX dataset being of a sufficient scale for judicious selection of training cohorts and effective implementation of deep learning methods. Much of the framework to interact with this data is automated in our pipeline, which allows for rapid model prototyping for a wide range of diseases and outcomes. In addition to making this a fully data-driven framework, the degree of automated preprocessing in our pipeline reduces the barrier of entry for potential end users, including physicians, researchers, and industry professionals who may not have the technical expertise to wrangle large-scale data and implement deep learning and machine learning models.

## 4 CONCLUSIONS

In this study, we demonstrated the utility of integrating multimodal deep learning approaches with federated healthcare datasets. Our pipeline achieves robust discriminatory scores on multiple disease prediction tasks, and given refinement and expansion, this methodology represents a promising step towards democratization of deep learning and artificial intelligence in a healthcare setting.

## A APPENDICES

### A.1 Introduction

### A.2 Materials and Methods

#### A.2.1 Supplementary Code2Vec Information

**Appendix Table 1:**
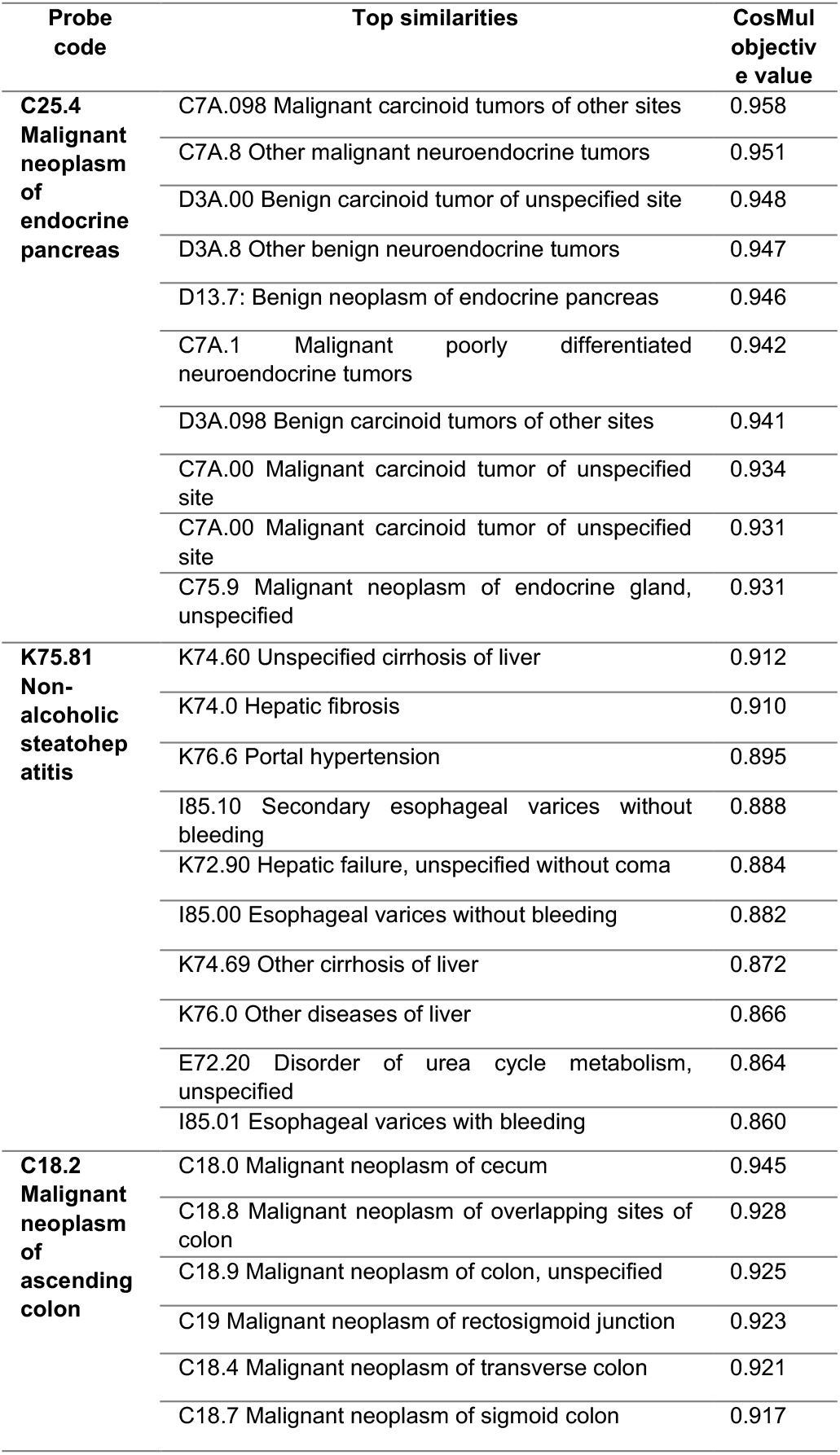
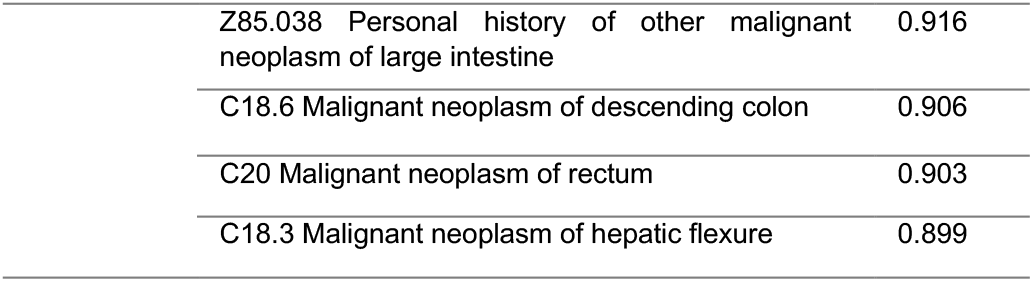
Selection of three probe ICD codes and their top 10 most similar codes, as evaluated with the cosine multiplication objective proposed by Levy and Goldberg[9].

The Code2Vec pipeline draws inspiration from several works for representation of code-based data. The Word2Vec algorithm, proposed by Mikolov et. al. [17], is perhaps the most well-known of a family of shallow neural networks that generate high-dimensional concept representations based on co-occurrence. The following alterations were made to the Word2Vec algorithm to make it suitable for representation of ICD and procedure codes– Training examples (or patient ‘sentences’) were generated by considering all unique diagnoses/procedures in a patient’s recorded medical history. The window size for the Word2Vec algorithm was extended to the maximum number of unique diagnoses encountered in a single patient’s history, and the specific model architecture used was skip-gram, which displays slightly better capture of semantic similarity versus the continuous bag-of-words (CBOW) architecture [16][17],.

Choi et al. proposed a simple but effective method of generating an overall patient representation from individual codes in their Med2Vec framework, which is to take a component-wise average of the vector representations of the codes [4]. We confirm the robustness of this approach (see discussion). In the embedding space, the similarity between vectors serves as an indicator of their semantic relatedness. Examples are given to show that the learned embeddings faithfully capture clinical relatedness in the case of diagnostic codes (**Appendix Table 1**) and the resulting high-dimensional representations of the codes can be dimensionally reduced to facilitate visualization.

### A.3 Results and Discussion

### A.4 Conclusions

### A.5 References

## ACKNOWLEDGMENTS

We appreciate the thoughtful insight and support from Dr. Jennifer Emond (Department of Epidemiology, Dartmouth College Geisel School of Medicine) and the Clinical Sciences team at TriNetX, Inc., particularly Seth Kuranz, Jennifer Stacy, Rutendo Kashwamba, Josh Hartman, and Sierra Luciano.

